# Nicotinic control of dopamine release shapes striatal output and engages adaptive action selection

**DOI:** 10.64898/2026.10.12.758519

**Authors:** Elif Beyza Guven, Samet Kocaturk, Michael W. Shiflett, Maxime Assous

## Abstract

Acetylcholine and dopamine significantly modulate striatal circuits and are essential for reward learning and action selection. Cholinergic interneurons can locally trigger dopamine release through β2-containing nicotinic acetylcholine receptors expressed on dopamine axons^1–3^. However, the causal contribution of this local cholinergic control of dopamine to behaviour and striatal output remains unresolved, with recent studies reaching contrasting conclusions about the extent and behavioural relevance of local acetylcholine–dopamine coupling^4–6^. Here, to directly test the causal contribution of this pathway, we conditionally deleted β2-containing nicotinic acetylcholine receptors from midbrain neurons in adult mice. This manipulation selectively abolished cholinergic interneuron-evoked dopamine release in the striatum. In a reward-based instrumental task, mice lacking this pathway showed decrements in effort that depended on the reinforcement context, and failed to adjust actions in a choice task when effort and reward costs varied. Substantia nigra β2-containing nicotinic acetylcholine receptor deletion also markedly weakened cholinergic interneuron-mediated disynaptic inhibition of striatal projection neurons, showing that disruption of β2-dependent cholinergic control of dopamine alters how cholinergic activity is transmitted to striatal output. Together, these findings establish the importance of local acetylcholine–dopamine coupling for regulating striatal output and influencing whether actions are sustained and adjusted as effort, reward, and outcomes change.

## Introduction

Adaptive behaviour requires animals to weigh the benefits of an action against the effort required to obtain those benefits, and to adjust behaviour when these relationships change. Dopamine (DA) is central to this process, contributing to behavioural activation, reinforcement learning, and the willingness to work for valued outcomes^7–13^. How DA signalling contributes to these different components remains incompletely understood.

Afferent neurons from the substantia nigra (SN) provide DA to the striatum, the main input structure of the basal ganglia. Importantly, the DA signal available to striatal circuits is determined not only by the firing of midbrain DA neurons^14^, but can also be regulated locally at dopaminergic axons. Acetylcholine (ACh) released from striatal cholinergic interneurons (CINs) provides a major mechanism for this local regulation. β2-containing nicotinic acetylcholine receptors (β2*-nAChRs) expressed on DA axons strongly regulate striatal DA release^15–18^. Synchronous CIN activation can evoke DA release through these receptors independently of action potentials initiated at DA neuron somata^1,2^, while direct axonal recordings have shown that nicotinic input can depolarise DA axons and initiate local action potentials^3,19^. More recent work has shown that nicotinic depolarisation can also transiently alter the responsiveness of DA axons to subsequent activity^20^. β2*-nAChRs therefore provide a powerful mechanism through which local cholinergic activity can dynamically shape dopaminergic output.

Despite the detailed cellular understanding of these CIN-DA interactions, the contribution of local cholinergic control of DA to behaviour remains less clear. CIN activity changes around salient events, rewards, and shifts in behavioural contingencies^21–24^, and perturbing striatal cholinergic signalling can impair adaptation when action-outcome relationships change^25–27^. Simultaneous measurements of ACh and DA in behaving animals have revealed highly dynamic relationships, including correlated, anticorrelated and temporally offset signals that vary with behavioural state, reward history and task demands^4,5,28,29^. These observations have led to contrasting views of the extent to which ACh directly controls DA release in vivo. Recent work has further linked local cholinergic regulation of DA to effortful behaviour with local nicotinic receptor blockade reducing high-effort responding^6^. Because β2*-nAChRs are also expressed by striatal GABAergic interneurons and on other afferents, including glutamatergic, serotonergic and long-range GABAergic afferents, local pharmacological manipulations lack cellular specificity^30–35^. The causal contribution of β2-nAChR-dependent cholinergic control of midbrain DA afferents to behaviour and striatal output therefore remains unresolved.

Here, we directly tested this mechanism by conditionally deleting β2-nAChRs from substantia nigra (SN) neurons in adult mice and examining the consequences for striatal DA signalling, behaviour, and local circuit function. This manipulation selectively abolished CIN- evoked DA release while preserving β2-independent dopamine release. Across complementary behavioural assays, β2 deletion reduced sustained responding as effort requirements increased, disrupted the normal experience-dependent reallocation of behaviour when greater reward required greater effort, and slowed the suppression of responding following reward omission. At the circuit level, β2 deletion also markedly weakened CIN-mediated disynaptic inhibition of spiny projection neurons (SPNs)^32,34,36^, revealing that loss of β2-dependent cholinergic control of midbrain signalling alters how cholinergic activity is transmitted to striatal output. Together, these findings identify nicotinic control of dopamine release as a causal regulator of striatal output and adaptive action regulation.

## Results

### Adult SN β2-nAChRs deletion selectively abolishes CIN-evoked dopamine release

CINs evoke striatal DA release through activation of β2-nAChRs expressed by nigrostriatal DA axons. To selectively disrupt this pathway in adulthood, we crossed ChAT-ChR2-eYFP mice with Chrnb2-floxed mice and generated a conditional substantia nigra β2 knockout (SN-β2-KO) by unilaterally injecting AAV2-hSyn-Cre-mCherry into the SN, while the contralateral hemisphere received control AAV2-hSyn-mCherry (Fig. 1a,b).

**Fig. 1.**
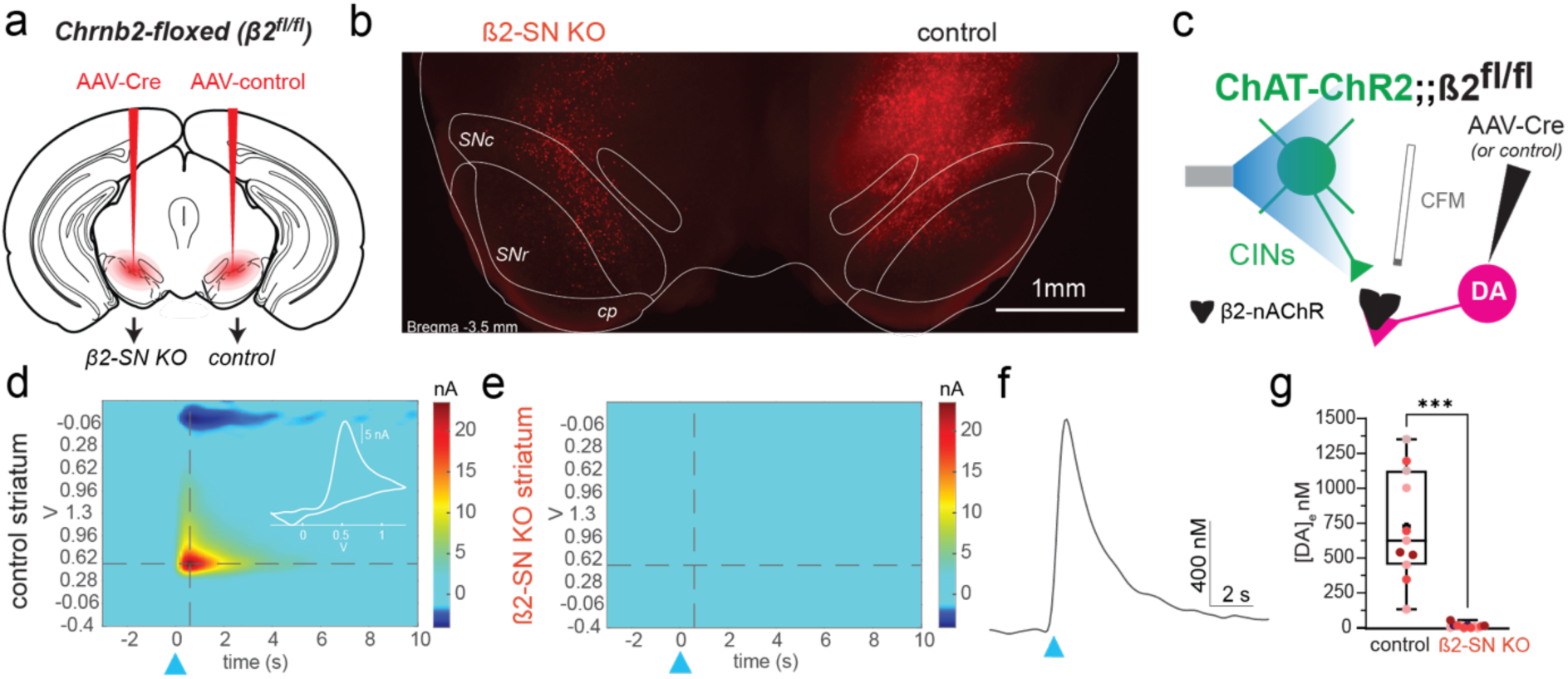
Conditional deletion of β2-containing nicotinic acetylcholine receptors from substantia nigra neurons abolishes cholinergic interneuron-evoked dopamine release. (a) Experimental strategy for conditional deletion of β2-nAChRs in the substantia nigra. Chrnb2-floxed mice received unilateral AAV2-hSyn-Cre-mCherry and contralateral AAV2-hSyn-mCherry injections. (b) Representative coronal section showing viral expression in the β2-SN KO and control hemispheres, with SNc, SNr and cerebral peduncle (cp) outlined. Scale bar, 1 mm. (c) Experimental configuration. Chrnb2-floxed mice were crossed with ChAT-ChR2-eYFP mice, and striatal dopamine release evoked by optogenetic activation of cholinergic interneurons (CINs) was measured using fast-scan cyclic voltammetry (FSCV). (d-e) Representative FSCV pseudocolour plots from control (d) and β2-SN KO (e) striata following optical stimulation of CINs (blue triangle). (f) Representative dopamine concentration transient evoked by CIN stimulation in control striatum. (g) Peak extracellular dopamine concentration evoked by CIN stimulation in control and β2-SN KO striata. Box plots show the median, interquartile range and minimum-to-maximum values; points represent individual measurements, with points of the same colour obtained from the same mouse. ***P < 0.0001, unpaired two-tailed t-test.

To determine whether this strategy efficiently abolished cholinergically evoked DA release, we performed fast-scan cyclic voltammetry (FSCV) in acute dorsal striatal slices while optogenetically activating CINs (2ms, 450nm, Fig. 1c). In control hemispheres, activation of CINs reliably evoked DA transients (peak amplitude: 170.9 ± 28.7 nM, *n* = 12 from N=4 mice), whereas no detectable DA release was observed in the contralateral SN-β2-KO hemisphere (6.4 ± 3.2 nM, *n* = 11 from the same N=4 mice; two-tailed unpaired t-test, p<0.0001; Fig. 1d-g).

To determine whether this manipulation selectively disrupted cholinergically evoked DA release rather than producing a general impairment, we next compared optogenetically and electrically evoked DA release (Extended Data Fig. 1). In control hemispheres, electrical stimulation elicited DA transients comparable in amplitude to those evoked by optogenetic activation of CINs (714.3 ± 68.9 nM, n=8 vs 671.9 ± 21.5 nM, n=6, two-tailed Welch’s t-test, p=0.572, Extended Data Fig. 1c-d). Consistent with previous studies, bath application of the β2-nAChR antagonist DHβE significantly reduced electrically evoked DA release (271.18 ± 70.02 nM, *n* = 4, two-tailed Welch’s t-test, p=0.0018, Extended Data Fig. 1e). Importantly, electrically evoked DA release in the SN-β2-KO hemisphere (340.17 ± 13.77 nM, *n* = 8) was not significantly different from DHβE-treated control slices (two-tailed Welch’s t-test, p=0.4002, Extended Data Fig. 1f-h). These findings indicate that SN-β2 deletion removes the β2-dependent cholinergic component of DA release while preserving β2-independent dopamine signalling. This model therefore provides a selective mean to test the physiological and behavioural consequences of disrupting CIN-evoked dopamine release..

### SN β2-nAChRs sustain instrumental responding under increasing effort requirements

We next asked whether disrupting nicotinic control of CIN-evoked dopamine release alters the ability of animals to sustain instrumental responding under conditions of increasing effort for reinforcement. β2-floxed mice received bilateral injections of either AAV2-hSyn-Cre-mCherry (SN-β2-KO) or AAV2-hSyn-mCherry (control) into the SN (Fig. 2a,b). Atlas-based reconstruction of all injection sites across the behavioural cohort confirmed targeting with comparable viral spread observed in control (n = 12) and SN-β2-KO (n = 15) mice (Extended Data Fig. 2). Mice were first trained in an operant task to perform a lever press to gain a food pellet. SN-β2-KO and controls showed no difference in acquisition of the lever press response (Extended Data Fig. 3a).

**Fig. 2.**
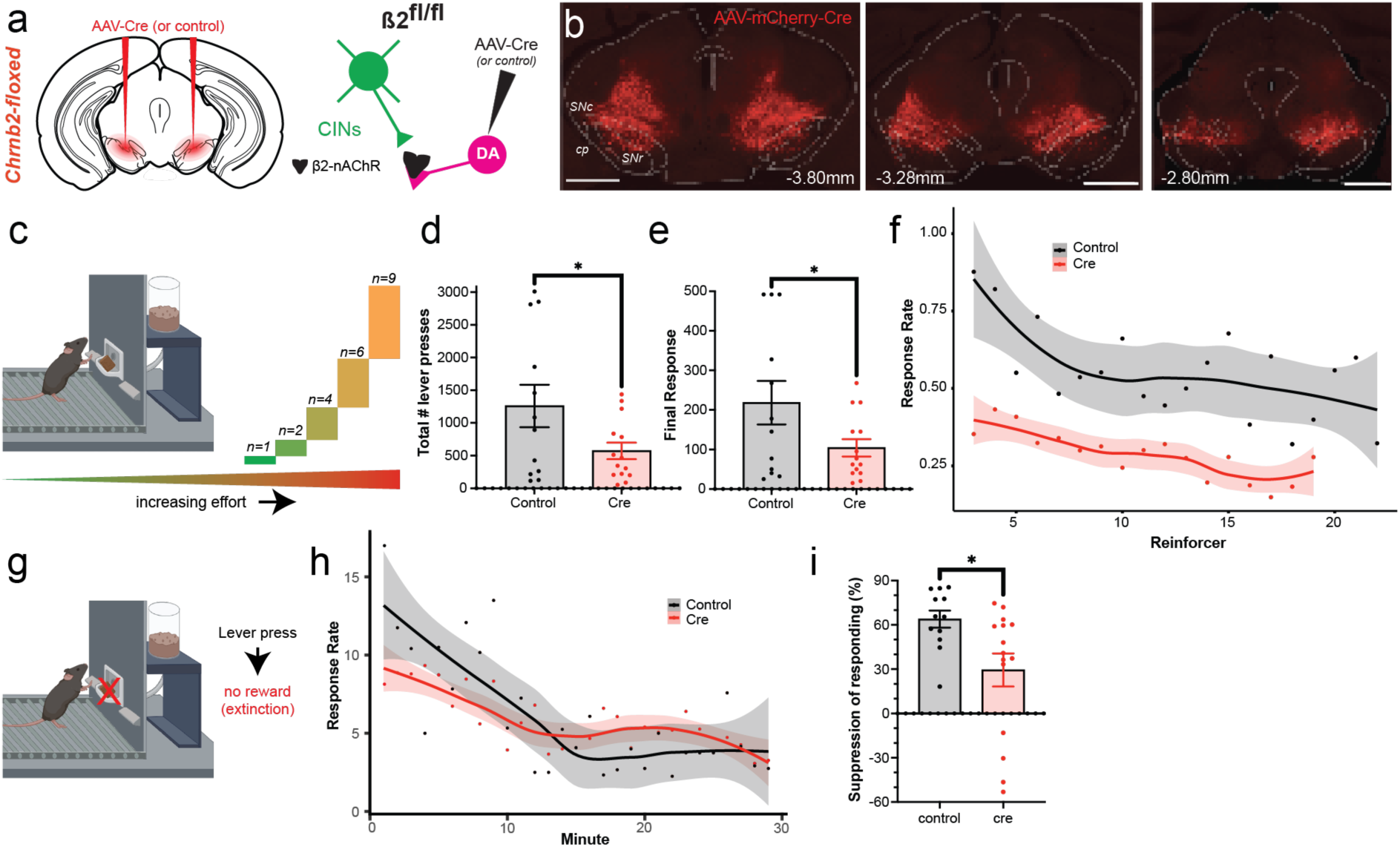
Nigrostriatal β2-containing nicotinic acetylcholine receptors promote sustained instrumental responding. (a) Experimental design illustrating unilateral deletion of β2-nAChRs from SN neurons by injection of AAV2-hSyn-Cre-mCherry. Control mice received AAV2-hSyn-mCherry injections. (b) Representative coronal sections illustrating mCherry expression. (c) Schematic of the progressive ratio task. Mice responded for food rewards under a progressive ratio schedule in which the number of lever presses required to obtain each successive reward increased until animals ceased responding. (d) Quantification of the total number of lever presses performed during the progressive ratio session in control and SN β2 knockout mice (Cre). (e) Quantification of the final response requirement completed before animals ceased responding. (f) Mean response rate across reinforcer deliveries with fitted curves for each group. Ribbon represents 95% confidence interval. (g) Schematic of the extinction task. Mice responded on the previously rewarded lever, with no outcome deliveries (h) Mean response rate across one-min bins of the extinction session, with fitted curves for each group. Ribbon represents 95% confidence interval. (i) Change in repsonse rate in the last 10 min of the session compared to the first 10 min. data points in (d), (e) and (i) are presented as individual animals with bar graphs showing mean ± SEM. Statistical comparisons were performed using unpaired t-tests. *p < 0.05.

We then tested mice on a progressive ratio task in which the number of lever presses required to obtain successive food rewards increased according to an exponential schedule (Rn = round (5e^0.2n^ - 5), Fig. 2c). Total lever presses in SN-β2-knockout mice were significantly reduced compared with control mice (Control: 1259 ± 325, SN-β2-KO: 571.8 ± 126.2; *P* = 0.0341, unpaired Welch’s t-test; Fig. 2d), indicating that disruption of SN β2-nAChRs reduced sustained instrumental responding as effort requirements progressively increased. Consistent with this observation, SN-β2-KO mice also terminated their session at a significantly lower response requirement than control animals (Control: 218.25 ± 54.94, SN-β2-KO: 104.07 ± 21.88; *P* = 0.0367, unpaired Welch’s t-test; Fig. 2e), demonstrating that SN-β2-KO mice disengaged from the task as effort requirements increased. Breakpoint showed a similar downward trend following SN-β2-KO (Control: 16.75 ± 1.55, SN β2-KO: 14 ± 1.08; *P* = 0.0805, unpaired Welch’s t-test; Extended Data Fig. 3b). To visualize behavioural persistence throughout the progressive ratio schedule, we plotted the fraction of animals remaining engaged as response requirements increased (Extended Data Fig. 3c). Whereas control mice continued responding across progressively higher ratio requirements, SN β2 knockout mice disengaged earlier, resulting in a leftward shift of the persistence curve. This is consistent with the reductions in total lever presses and final completed response requirement observed in SN β2 knockout mice.

To examine how the instrumental response rate changed over the session, we fitted the following exponential decay function to group mean response rates: *y=a*exp(x*-b)* with *y* being the response rate and *x* being the trial number (Fig. 2f). The peak response rate (*a*) and decay rate (*b*) were estimated for each animal independently and group level statistics performed on these values. SN-β2-KO mice showed a significantly lower peak response rate (*a*) (P=0.001) (Extended Data Fig. 3d). The decay rate (*b*) did not differ by group (P=0.22, Extended Data Fig. 3d). Together, these findings indicate that nicotinic control of dopamine release contributes to sustained instrumental responding in the progressive ratio task.

### Deletion of SN β2-nAChRs impairs extinction of previously reinforced responding

We next examined whether disrupting nicotinic control of dopamine release alters the ability to disengage from a previously reinforced action when reinforcement was no longer available. Mice were given additional instrumental training and then underwent a 30-min extinction session during which lever presses no longer resulted in reward delivery (Fig. 2g). Both groups progressively reduced responding across the session, demonstrating extinction of the previously reinforced response. However, the temporal profile of extinction differed significantly between genotypes. Using the same curve-fitting function as described above, we found that the peak response rate (*a*) and decay rate (*b*) were significantly altered following SN β2 deletion (Fig. 2h). SN-β2 knockout mice showed a significantly lower peak response rate (*a*) (P=0.018) and a significantly lower decay rate (*b*) (P=0.04) (Extended Data Fig. 3e). To quantify the degree to which individual animals suppressed responding over the course of extinction, we compared responding during the first 10 min with that during the final 10 min of the session. Control mice reduced their responding by 63.9%, whereas SN-β2-KO mice showed only 29.5% suppression, indicating substantially weaker disengagement from the previously rewarded action (Fig. 2i; Welch’s t(20.65) = 2.731, P = 0.0126). This difference was not explained by greater overall instrumental activity, as the total number of lever presses emitted across the 30-min extinction session was similar between control and SN β2-KO mice (Extended Data Fig. 3f, Control: 193.9 ± 29.6 presses; SN-β2-KO: 178.7 ± 24.6 presses; Welch’s t(22.90) = 0.394, P = 0.6969).

Thus, SN β2 deletion did not alter the overall amount of responding during extinction but selectively reduced the vigour and increased the persistence of responding over the session, consistent with reduced behavioural disengagement when the learned action– outcome relationship was no longer reinforced.

### SN β2-nAChRs support adaptive choice under reward–effort conflict

We next asked whether this pathway contributes to decisions requiring integration of effort cost and reward value. Control and SN-β2-KO mice were tested in an effort-based decision-making task in which they chose between one lever operating under a fixed effort requirement (FR-5 schedule) and a second lever whose effort requirement varied (VR schedule, Fig. 3a).

**Fig. 3.**
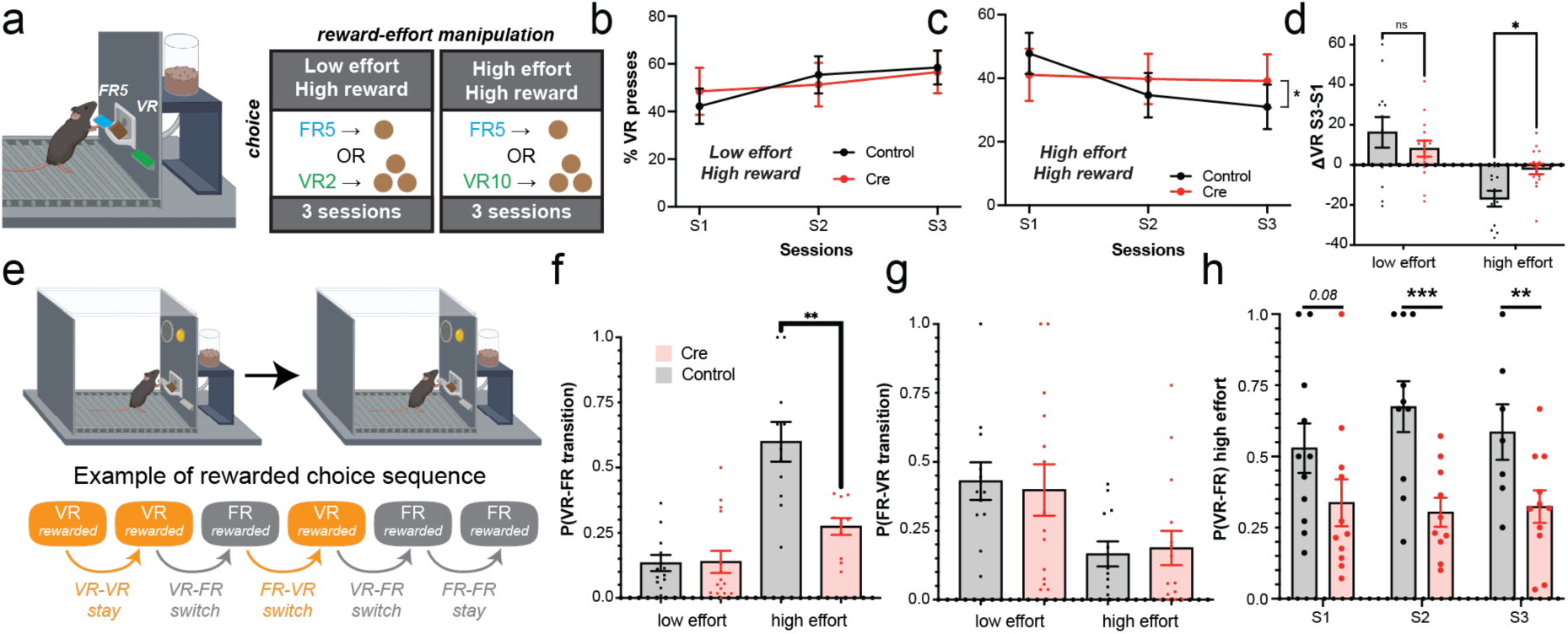
SN β2-nAChRs are required for adaptive effort-based decision making under high effort–high reward conflict. (a) Choice task in which mice selected between a fixed-ratio option delivering one pellet and a variable-ratio option delivering three pellets under low- or high-effort conditions. (b,c) Percentage of responses allocated to the variable-ratio option across three sessions in the low-effort (b) and high-effort (c) conditions. (d) Change in variable-ratio choice from session 1 to session 3. (e) Example rewarded choice sequence illustrating stay and switch transitions between FR and VR responses. (f,g) Probability of VR→FR (f) and FR→VR (g) transitions under low-and high-effort conditions. (h) Probability of VR→FR transitions across sessions in the high-effort condition. Points represent individual animals; bars show mean ± s.e.m.

Additionally, responses on the VR lever delivered 3 food pellets, compared to a single food pellet following responses on the FR lever. Under these conditions, mice had to account for differences in both effort cost and reward value when selecting a response. When the larger reward was associated with the lower-effort VR2 option, both control and SN-β2-KO mice showed a similar adaptive increase in their proportion of VR responding across sessions (session: F2.00,49.88 =4.43, P=0.0170), with no effect of genotype (P=0.9979) or session × genotype interaction (P=0.4286; Fig. 3b). In contrast, when obtaining the larger reward required greater effort, SN-β2-KO showed no adaptive decrease in VR lever preference (Fig. 3c). Unlike control mice, which progressively reduced their preference for the high-effort/high-reward VR option, with significant differences between S1 and S2 (P=0.0024) and between S1 and S3 (P=0.0039), SN-β2-KO mice showed no corresponding change between S1 and S2 (P=0.9904) or S1 and S3 (P=0.8522). There was a significant effect of session (F1.83,45.66 =7.24, P=0.0024) and, critically, a significant session × genotype interaction (F1.83,45.66 =4.60, P=0.0176). Thus, whereas control mice progressively reallocated their choices away from the costly VR option with repeated experience, this adaptation was markedly attenuated following SN β2 deletion.

To quantify the net change in choice across repeated sessions, we calculated the difference in VR preference between Sessions 1 and 3 for low and high effort conditions when rewards were unequal (Fig. 3d). Adaptation scores varied by group and genotype, with a significant group x genotype interaction (F1,50 = 6.04, P=0.0174). Planned comparisons revealed a difference between genotypes specifically under the high effort condition: controls reduced VR preference by 16.82% compared with only 1.98% in SN-β2-KO mice (Šídák-adjusted P=0.0267); no genotype differences were detected in the low effort condition.

Importantly, during the first session of each condition, VR preference did not differ according to condition (P=0.8892) or genotype (P=0.7101), with no condition × genotype interaction (P=0.3717) and no genotype difference within any individual (all P≥0.7961; Fig. Extended Data Fig. 5a). Thus, the lack of adaptive responding under the high effort condition in SN-β2-KO mice was not attributable to a pre-existing difference in choice at the beginning of testing. Moreover, the lever response rate during the high effort condition did not differ between genotypes (control mean = 8.93 ± 3.99 SD; SN-β2-KO = 8.41 ±4.09 SD, Extended Data Fig. 5b), suggesting the effects on choice were not influenced by sluggish responding in SN-β2-KO mice. Likewise, the average latency between pellet retrieval and first lever response did not differ between groups (control = 15.29 s ± 22.63 SD; SN-β2-KO = 19.21± 36.60 SD). The selective change in choice allocation therefore occurred without evidence of a gross reduction in instrumental responding.

The inability of SN-β2-KO mice to disengage from costly but richly rewarded actions was not due to a deficit in behavioural flexibility generally. When mice experienced the same reward size on both levers (1 food pellet; Extended Data Fig. 4a-b), both control and SN β2-KO mice adjusted their choices according to the relative effort associated with the VR option (Extended Data Fig. 4c-d). When the VR lever required less effort than the FR5 alternative, VR preference increased across sessions (session: F1.83,45.81=14.98, P<0.0001), with no effect of genotype (P=0.7177) or session × genotype interaction (P=0.7594). When both options imposed comparable effort requirements, preference remained relatively stable, with no significant effect of session (P=0.5342), genotype (P=0.5871, or session × genotype interaction (P=0.6944). Conversely, when the VR option required greater effort, animals progressively reduced their preference for this option (session: F1.90,47.51=14.83, P<0.0001), again without a significant genotype effect (P=0.3433) or session × genotype interaction (P=0.6835). Thus, under equal reward conditions, mice adapted their behaviour to changing effort requirements, with no behavioural impairment produced by β2-nAChR deletion.

Together, these findings indicate that nigrostriatal β2-nAChRs are important when animals must adapt their choices under competing reward and effort demands.

### SN β2-nAChRs deletion selectively reduces disengagement from a rewarded high-effort/high-reward option

We next examined the trial-by-trial organization of choice when mice had to consider differences in both reward magnitude and effort, to determine what behavioural strategy accompanied the impaired adaptation observed during testing under the high reward condition. Each rewarded VR choice was classified according to whether the subsequent completed rewarded choice remained on the VR option or transitioned to FR (Fig. 3e). The same approach was used to quantify transitions in the opposite direction.

The probability of transitioning from VR to FR varied markedly depending on the effort/reward contingency and genotype. Overall, there was a significant effort condition × genotype interaction (F1,20 =28.56, P<0.0001). In the high effort condition, control mice showed a substantially greater probability of switching from VR to FR than SN-β2-KO mice (Control: 0.599; SN-β2-KO: 0.274; P=0.0081) (Fig 3f). In contrast, VR→FR switching did not differ between genotypes in the low effort high reward condition (P = 0.81). The genotype effect on transitions away from the VR option was therefore specifically expressed when that option combined a larger reward with a greater effort requirement.

This alteration in choice was also directionally selective. FR→VR transition probabilities varied substantially between high and low effort conditions (condition: F1,24 =32.04, P<0.0001; Fig. 3g) but were entirely unaffected by SN-β2-KO, with neither an overall genotype effect (F1,25 =0.007, P=0.9326) nor a condition × genotype interaction (F1,24 =0.093, P=0.7625). Accordingly, none of the genotype comparisons within individual conditions was significant, including in the high effort condition (Control: 0.166; SN β2-KO: 0.187; P=0.9995). Thus, SN-β2-KO mice were not generally less likely to switch between alternatives; rather, the deficit was specific to switching away from the rewarded high-effort option.

To examine the temporal organization of this effect, we next resolved VR→FR switching across the three high effort sessions (Fig. 3h). Mixed-effects analysis revealed a significant overall effect of genotype (F1,20 =12.61, P=0.0020), without a significant effect of session (F2,34 =0.92, P=0.4085) or session × genotype interaction (F2,34 =1.65, P=0.2067). Between-group comparisons showed lower VR→FR switching in SN-β2-KO mice during S2 (P=0.0010) and S3 (P=0.0039), whereas the difference during S1 did not reach significance (P=0.0803). These findings demonstrate a robust overall reduction in the tendency of SN-β2-KO mice to disengage from the VR option during the high effort test condition.

These findings suggest that β2-nAChR-dependent regulation of dopamine release contributes to the adaptive disengagement from high-value actions when their associated effort costs must be considered.

### SN β2-nAChRs support CIN-mediated control of striatal output

The behavioural effects of midbrain β2-nAChR deletion suggested that this pathway may influence how cholinergic signals are translated into striatal output. CINs exert rapid control over SPNs through a disynaptic GABAergic circuit involving β2-nAChR-dependent recruitment of striatal interneurons, a pathway proposed to transmit behaviourally relevant cholinergic signals to projection neurons^32–34,36–39^. We therefore asked whether disrupting β2-dependent signalling in midbrain afferents alters this CIN-driven control of SPNs.

Whole-cell voltage-clamp recordings were obtained from SPNs while CINs were optogenetically stimulated using ChAT-ChR2-eYFP;;β2-floxed mice (Fig. 4a). In control hemispheres, brief blue-light stimulation evoked the characteristic biphasic IPSC comprising kinetically distinct fast and slow components (Fig. 4b). In contrast, SN-β2-KO markedly attenuated both components (Fig. 4c). Quantification revealed a profound reduction in the peak amplitude of the fast IPSC following SN β2 knockout (Control: 482.6 ± 103.1 pA, SN-β2-KO: 77.2 ± 22.5 pA, Mann–Whitney, *P* < 0.0001; n = 19 control and 22 SN-β2-KO recordings obtained from N=4 mice; Fig. 4d). Likewise, the slow IPSC component was significantly reduced in SN β2-KO hemispheres (Control: 459.9 ± 85.5 pA, SN β2-KO: 95.4 ± 25.4 pA, Mann–Whitney, *P* < 0.0001; Fig. 4e). Together, these results demonstrate that β2-containing nicotinic receptors expressed by nigrostriatal dopamine neurons are required for the full expression of CIN-mediated disynaptic inhibition of SPNs.

**Fig. 4.**
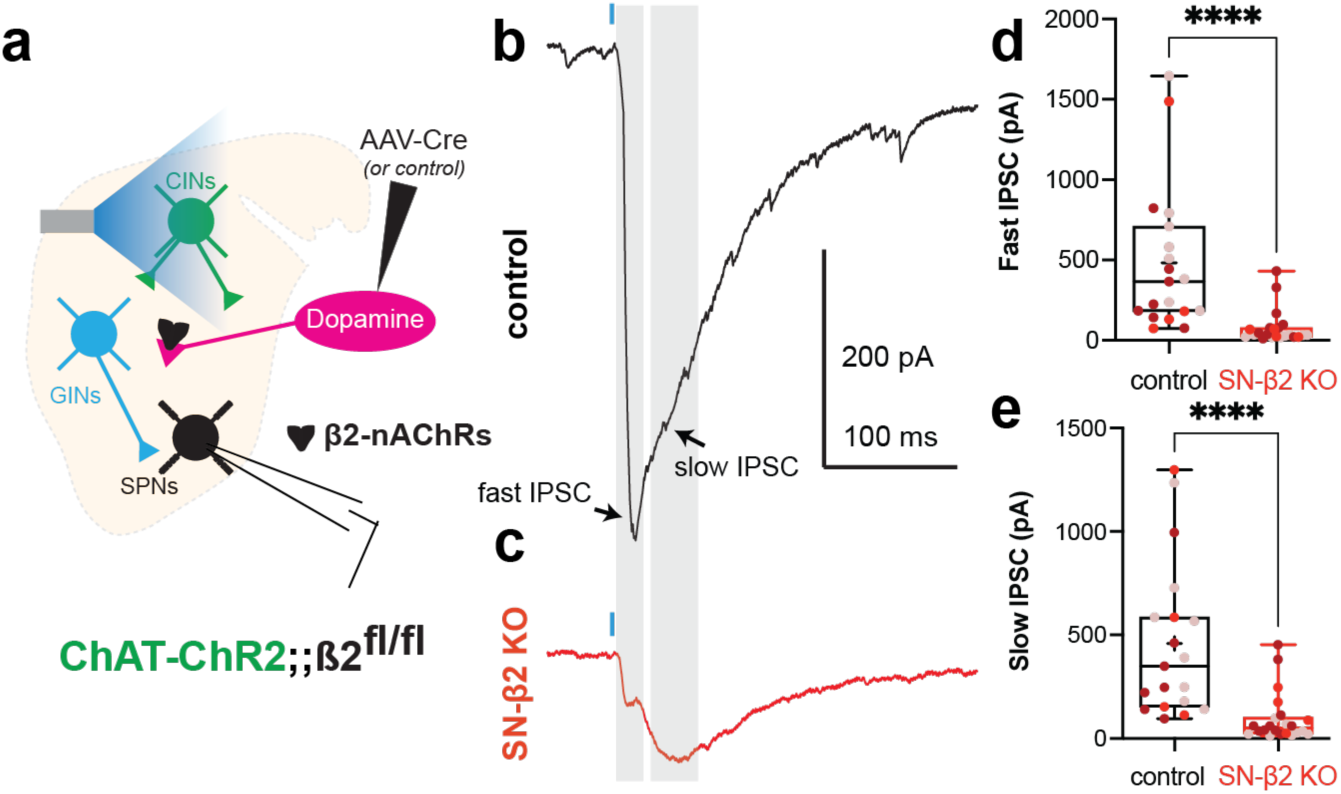
Midbrain β2-nAChRs regulate CIN-mediated disynaptic inhibition of SPNs. (a) Experimental schematic for whole-cell voltage-clamp recordings from SPNs in ChAT-ChR2;Chrnb2fl/fl mice following unilateral SN injection of AAV-hSyn-Cre-mCherry or control AAV. (c,d) Representative CIN-evoked IPSCs recorded from control (c) and β2-SN KO (d) hemispheres. (e,f) Peak amplitudes of fast (e) and slow (f) IPSC components. Points represent individual cells; points of the same colour indicate recordings from the same mouse. Box plots show median, interquartile range and minimum-to-maximum values. ****P < 0.0001, two-tailed Mann–Whitney test.

We next asked whether acute disruption of dopamine receptor signalling influenced the strength of this circuit. Combined blockade of D1 and D2 receptors significantly reduced both components of the CIN-evoked IPSC (Extended Data Fig. 6a,b; fast IPSC: Control, 753.5 ± 152.9 pA; SCH-23390 + sulpiride, 250.9 ± 85.69 pA; Wilcoxon matched-pairs test, *P* = 0.0312, n = 6; slow IPSC: Control, 483.4 ± 116.6 pA; SCH-23390 + sulpiride, 207.2 ± 107.6 pA; *P* = 0.03). These findings indicate that ongoing dopaminergic signalling contributes substantially to the strength of CIN-mediated inhibition. Consistent with this circuit being sensitive to the integrity of nigrostriatal dopamine signalling, CIN-mediated inhibition of SPNs is also profoundly disrupted following 6-OHDA lesioning^38,39^ or the MCI-Park model^38^. Thus, disrupting CIN-driven midbrain signalling alters not only local DA release but also a major route through which cholinergic activity controls striatal output. These results identify a downstream circuit consequence of disrupting β2-dependent CIN–midbrain signalling that is positioned to alter how behaviourally relevant cholinergic activity is transmitted to striatal output neurons.

## Discussion

Our findings identify β2-nAChR-dependent cholinergic control of striatal DA release as an important regulator of adaptive behaviour. Adult midbrain β2 deletion abolished CIN-evoked DA release, reduced sustained responding when effort requirements and reinforcement schedules change, and altered experience-dependent allocation between actions differing in effort and reward. These behavioural effects were accompanied by a marked reduction in CIN-mediated disynaptic control of SPNs, indicating that loss of β2-dependent cholinergic control of midbrain afferents is associated not only with loss of CIN-evoked dopamine release, but also with altered cholinergic regulation of striatal output. Together, these findings suggest a complex role for SN-β2 signalling in determining whether to sustain or reallocate actions as effort costs and consequences change

β2-nAChRs provide a rapid route through which CINs regulate striatal DA axons. Synchronous CIN activity can evoke DA release independently of somatic firing, while nicotinic activation can also shape the subsequent responsiveness of DA axons to ongoing activity^1–3,19,20^. We interrogated this pathway using an adult, SN-centred viral deletion of β2-nAChRs. This strategy provides greater cellular specificity than local nicotinic antagonism, which also affects β2-containing receptors on other striatal elements^6,31–35,37,38^. Furthermore, the adult deletion limits the opportunity for developmental adaptation associated with constitutive or early developmental deletion^4,40^. Although our manipulation was not restricted to DA neurons, it abolished CIN-evoked DA release, confirming effective disruption of this pathway. Consistent with a selective loss of nicotinic control, rather than a generalized impairment of DA release, electrically evoked DA signals were comparable between control and SN-β2 knockout mice in the presence of the β2-nAChRs antagonist DHβE, indicating that β2-independent components of DA release remained intact. Together, these results validate the use of this viral strategy to examine how nicotinic control of DA release influences striatal circuit function and behaviour.

To assess SN-β2’s role in effortful behaviour, mice performed a free-operant task with food reinforcers delivered under a progressive ratio schedule or withheld entirely in an extinction task. In both tasks, the peak response rate was significantly lower in SN-β2 knockout mice, suggesting reduced effortful task engagement. Additionally, in the progressive ratio task, SN-β2 knockout mice made fewer lever presses and achieved a lower final response ratio, despite normal acquisition of instrumental responding. Because progressive-ratio performance also incorporates increasing delays to reward, pausing and eventual stopping^41^, we interpret this phenotype as reduced sustained instrumental engagement rather than a selective deficit in effort valuation or motivation. Additionally, these findings converge with recent work showing that cholinergic regulation of dopamine becomes increasingly important as effort demands rise. Touponse et al. found that higher effort requirements enhance nicotinic amplification of reward-evoked dopamine and that disrupting this mechanism compromises high-effort responding^6^. However, in the extinction task, we found that SN-β2 mice showed greater persistence in responding in the absence of reinforcement. This finding suggests a complex role for SN-β2 signalling with respect to effort, with contributions to both sustaining and reducing effort depending on the reinforcement context.

Further illustrating the complex relationship between SN-β2 signalling and effort, β2 deletion altered how choice allocation evolved across experience when mice weighed options that differed in both effort and reward size. Unlike control mice that shifted their responses toward lower-effort options, SN-β2-knockout mice persisted in choosing options that deliver larger rewards, despite their higher effort costs. The genotype difference therefore reflected failure to develop the normal experience-dependent shift away from the costly option rather than increasing preference for rewards earned under high effort. Importantly, genotype differences were not observed when the alternatives were matched for reward value or when the higher-value option carried the lower effort cost, arguing against a generalized side bias or inability to discriminate differences in effort costs or reward magnitude. Instead, the phenotype emerged specifically when animals had to resolve a conflict between greater reward and greater effort. Dopamine is strongly implicated in effort–cost decision-making^9,11,42^. Yet broad disruption of mesolimbic dopamine generally biases behaviour away from costly alternatives, whereas β2 deletion here preserved selection of the high-cost option. This distinction underscores the idea that disrupting β2-dependent CIN control is not equivalent to globally reducing dopamine transmission but instead interferes with a temporally and spatially restricted mode of cholinergic regulation.

Perseveration of the high-effort/high-reward option in SN-β2 knockout mice, with normal transitions in the opposite direction, further argues against a generalized switching deficit and instead points to reduced disengagement from a costly but still valuable action. Striatal CINs are well positioned to influence such behavioural adjustment. Parafascicular input to CINs contributes to incorporating new action–outcome information, and manipulation of CIN activity alters set shifting and behavioural adjustment following changes in instrumental contingencies^25–27^. Extinction provided an independent convergence with this reduced disengagement: both groups reduced responding after reward omission, but SN-β2-deficient mice showed a slower decay in response rate and weaker suppression across the session. Interestingly, unlike the extinction task, the response decay rate in the progressive ratio task was unaffected in SN-β2 knockout mice. One possibility for this difference in decay rate between the tasks is that outcome delivery in the progressive ratio task was sufficient to enable mice to adjust their effort, but in the absence of incentive updating in the extinction task SN- β2 mice were unable to adjust their actions. Together, these observations suggest that β2- dependent cholinergic control of midbrain afferents contributes to the context-dependent adjustment of established actions.

The marked reduction in CIN-mediated disynaptic inhibition of SPNs provides a plausible circuit route through which loss of β2-dependent midbrain signalling could alter action control. Synchronous CIN activity recruits β2-nAChR-expressing GABAergic interneurons to generate fast and slow GABAergic responses in SPNs, providing a rapid pathway through which phasic cholinergic signals can influence striatal output^32–34,37,38^. Acute blockade of D1- and D2-dopamine receptors also reduced CIN-evoked disynaptic inhibition, indicating that dopamine receptor signalling contributes to the expression of this circuit. Such a mechanism is relevant to the organization of SPN population activity during action selection. Direct- and indirect-pathway SPNs (dSPNs and iSPNs, respectively) are co-recruited during behaviour but can make distinct contributions to the maintenance and updating of action policies^43–45^. In particular, iSPNs have been implicated in switching away from unsuccessful actions and updating action–outcome associations, whereas dSPN activity can favour repetition of selected or rewarded actions^44,45^. These functions are relevant to the selective reduction in VR→FR transitions observed here. However, our physiology does not indicate a pathway-specific circuit deficit: CIN-driven disynaptic responses were broadly reduced across recorded SPNs, and dopamine depletion similarly weakens this circuit in both dSPNs and iSPNs^38^. Rather than selectively affecting one output pathway, loss of β2-dependent cholinergic signalling may therefore disrupt the coordinated reconfiguration of SPN ensembles as behavioural demands and action values change.

This circuit is also well positioned to regulate behaviourally relevant cortical and thalamic input by shaping the temporal integration of glutamatergic excitation^38^. Recent studies show that thalamic activity can recruit CINs to generate nicotinic-dependent dopamine release in dorsomedial striatum (DMS), but not dorsolateral striatum (DLS), during learning, with this coupling varying with task engagement and recent dopamine history^46^. Frontal cortical inputs can similarly recruit CINs to drive cholinergic-dependent dopamine signals in DMS^47^. Thus, cortical and thalamic signals can converge on CINs to regulate both local dopamine release and inhibitory microcircuits controlling SPNs, providing a mechanism through which behaviourally relevant input could reshape striatal output in a region- and state-dependent manner.

β2-dependent CIN control of dopamine release is likely to interact with other cholinergic mechanisms that regulate SPN excitability and synaptic input^31,35,48,49^. Dopamine and acetylcholine jointly shape corticostriatal plasticity, with coordinated cholinergic pauses, phasic dopamine and SPN depolarization promoting potentiation of active corticostriatal synapses^50^. Loss of this β2-dependent dopamine signal could therefore alter both the immediate processing and longer-term strengthening of behaviourally relevant cortical and thalamic inputs, providing a potential mechanism for the abnormal adjustment of effortful actions observed here.

Overall, our findings identify a role for β2-nAChR-dependent cholinergic control of midbrain afferents in determining when effortful actions should be sustained, reallocated or abandoned. By linking behaviourally relevant cholinergic signals to local dopamine release and the regulation of SPN output, this pathway may help translate recent effort and reward experience into subsequent action choice. Disruption of this mechanism could therefore contribute to maladaptive persistence, abnormal effort allocation and impaired behavioural adjustment in disorders including Parkinson’s disease and addiction.

## Acknowledgements

We thank Fulva Shah for excellent technical assistance and expertise in maintaining the transgenic mouse colonies, and Michael Crair (Yale University) for providing the *Chrnb2*-floxed mice. We would like to thank Jonathan Ghaly and Daniel Marin for assistance with acquisition of behavior data. We also thank members of the Assous laboratory for helpful discussions and comments.

## Author contributions

S.K. and E.B.G. performed experiments, analysed data and contributed to experimental design. M.W.S. designed experiments, supervised research, analysed data, contributed to obtaining research funding and wrote the manuscript. M.A. conceived and designed the study, supervised the research, obtained funding, performed experiments, analysed and interpreted data, and wrote the manuscript.

## Data availability

The data supporting the findings of this study are available within the Article and its Extended Data. Additional data are available from the corresponding author upon reasonable request. Source data underlying the figures will be deposited in a public repository before publication.

## Code availability

Custom code used for behavioural analyses, curve fitting and physiological recordings is available from the corresponding author upon reasonable request.

## Additional information

Correspondence and requests for materials should be addressed to Maxime Assous.

## Methods

### Animals

Adult male and female mice were used for all experiments. For ex vivo experiments examining β2-containing nicotinic acetylcholine receptors (β2-nAChRs) expressed by nigrostriatal neurons, ChAT-ChR2-eYFP mice (Tg(Chat-COP4*H134R/eYFP, Slc18a3)6Gfng/J; Jackson Laboratory) were crossed with homozygous Chrnb2-floxed mice (β2-floxed, Chrnb2-fl/fl; Chrnb2-tm1.1Mccl; provided by Prof. Michael Crair, Yale University). In ChAT-ChR2-eYFP mice, channelrhodopsin-2 and eYFP are expressed under control of the ChAT promoter, enabling optogenetic activation and visualisation of striatal cholinergic interneurons (CINs). Conditional deletion of β2-nAChRs was achieved through Cre-mediated recombination of the floxed Chrnb2 allele.

ChAT-ChR2-eYFP::β2-floxed mice were used for fast-scan cyclic voltammetry (FSCV) and whole-cell electrophysiology experiments. Behavioural experiments were performed in Chrnb2-floxed mice without constitutive ChR2 expression.

Mice were 3–7 months of age. Animals were housed in groups of up to four per cage under a 12-h light/dark cycle with ad libitum access to food and water except during behavioural experiments requiring controlled food access.

All experiments were conducted in accordance with the National Institutes of Health Guide for the Care and Use of Laboratory Animals and were approved by the Rutgers University Institutional Animal Care and Use Committee. Procedures conducted in the United Kingdom were performed in compliance with the UK Animals (Scientific Procedures) Act 1986 under Home Office Licence PP7595333 and with the approval of the local Cardiff University Ethics Review Committee.

### Conditional deletion of β2-nAChRs from substantia nigra neurons

Mice were anaesthetised with isoflurane (1–2.5% in O2, 1 L/min) and positioned in a stereotaxic frame. Bupivacaine was administered subcutaneously at the incision site and a small craniotomy was made above each injection site.

The substantia nigra pars compacta (SNc) was targeted at AP −2.7 mm, ML ±1.35 mm relative to bregma and DV −4.2 mm relative to the brain surface. Viral vectors were delivered through pulled glass micropipettes with an approximately 30–40 μm inner tip diameter using a Nanoject II Injector (Drummond Scientific).

For ex vivo experiments, ChAT-ChR2-eYFP::β2-floxed mice received AAV2-hSyn-mCherry-Cre (UNC Vector Core, AV6145B) into the SNc of one hemisphere and AAV2-hSyn-mCherry (UNC Vector Core, AV5033E) into the contralateral SNc. A total volume of 990 nL was delivered per injection at 13.8 nL every 5 s. This within-animal design generated a SN β2-KO hemisphere and a corresponding control hemisphere in the same animal.

For behavioural experiments, homozygous β2-floxed mice received bilateral SNc injections of either AAV2-hSyn-mCherry-Cre (β2 SN knockout) or AAV2-hSyn-mCherry (control).

Following viral delivery, the micropipette was left in position for 10 min before being slowly withdrawn. Mice received ketoprofen and buprenorphine for postoperative analgesia and were allowed at least 4 weeks for viral expression before experimentation.

### Acute brain-slice preparation

Adult mice were anaesthetised with ketamine (80 mg/kg) and xylazine (20 mg/kg) and transcardially perfused with ice-cold N-methyl-D-glucamine (NMDG)-based cutting solution containing, in mM: 103 NMDG, 2.5 KCl, 1.2 NaH2PO4, 30 NaHCO3, 20 HEPES, 25 glucose, 101 HCl, 10 MgSO4, 2 thiourea, 3 sodium pyruvate, 12 N-acetyl cysteine and 0.5 CaCl2. The solution was continuously equilibrated with 95% O2/5% CO2 and adjusted to pH 7.2–7.4.

Brains were rapidly removed and 300-μm coronal sections containing the dorsal striatum were prepared using a VT1200S vibratome (Leica). Slices were incubated in oxygenated NMDG solution at 35°C for 5 min and subsequently transferred to standard artificial cerebrospinal fluid (aCSF) containing, in mM: 124 NaCl, 2.5 KCl, 1.2 NaH2PO4, 26 NaHCO3, 1 MgCl2, 2 CaCl2, 10 glucose and 3 sodium pyruvate. Slices were maintained at approximately 25°C until use. During recordings, slices were continuously perfused with oxygenated aCSF at 32–34°C at a flow rate of 2–4 mL/min.

### Whole-cell electrophysiology

SPNs were visualised using an Olympus BX50-WI microscope equipped with a 40× long-working-distance water-immersion objective and infrared differential-interference contrast optics. Patch electrodes had a resistance of 3–5 MΩ.

For voltage-clamp recordings of CIN-evoked inhibitory postsynaptic currents (IPSCs), patch electrodes contained a CsCl-based internal solution consisting of, in mM: 125 CsCl, 0.1 EGTA, 10 HEPES, 2 MgCl2, 4 Na2ATP and 0.4 Na2GTP. Biocytin was added to the internal solution to permit post hoc visualisation of recorded neurons. SPNs were voltage-clamped at −70 mV. Membrane currents were recorded using a MultiClamp 700B amplifier (Molecular Devices). Depending on the recording setup, data were digitised and acquired using either AxoGraph (v1.3.4) or Signal (Cambridge Electronic Design). Sampling rates were 20–40 kHz.

CIN-evoked IPSCs in SPNs contain kinetically separable fast and slow GABAA receptor-mediated components. Peak amplitudes of the fast and slow components were therefore quantified independently. Electrophysiological traces were analysed using AxoGraph, Signal and custom MATLAB scripts.

### Optogenetic stimulation of cholinergic interneurons

CINs were activated through constitutive ChR2-eYFP expression in ChAT-ChR2-eYFP::β2-floxed mice. Blue-light stimulation was delivered using a high-power LED (450 nm; LZC-R0H1000000, Mouser Electronics). CIN activation consisted of a single 2-ms blue-light pulse delivered at 30-s intervals.

This stimulation paradigm was used both to evoke striatal dopamine release for FSCV experiments and to recruit CIN-dependent disynaptic inhibition during SPN recordings.

### Fast-scan cyclic voltammetry

Extracellular dopamine release was measured using an NPI fast-scan cyclic voltammetry (FSCV) system or an Ensman EI-400 biopotentiostat (Cypress Systems) and digitized via an NI PCI-6052E PC board (National Instruments). Carbon-fibre electrodes had a fibre diameter of approximately 7–8 μm. A triangular voltage waveform from −0.4 V to +1.3 V and back to −0.4 V versus Ag/AgCl was applied at 400 V/s with a sampling frequency of 10 Hz. Dopamine was identified from its characteristic oxidation and reduction peaks. Carbon-fibre electrodes were post-calibrated in experimental medium containing 75, 100, 250 and 500 nM and 1 μM dopamine hydrochloride (Sigma). Dopamine release was additionally evoked by local electrical stimulation in the dorsal striatum. Responses obtained under control conditions and following β2-nAChR blockade with dihydro-β-erythroidine (DHβE; 1 μM) were compared with electrically evoked dopamine release in SN β2-KO striatum.

### Pharmacological manipulation of CIN-evoked inhibition

To determine the contribution of dopamine receptor signalling to CIN-evoked disynaptic inhibition of SPNs, D1- and D2-family receptors were blocked simultaneously with SCH-23390 (10 μM; Sigma-Aldrich) and sulpiride (5 μM; Tocris), respectively (bath applied in oxygenated aCSF). CIN-evoked fast and slow IPSCs were recorded in ChAT-ChR2-eYFP slices before and during drug application. The combined antagonist treatment was evaluated within individual neurons using paired measurements.

### Histology and imaging

For post hoc anatomical verification, mice were deeply anaesthetised with ketamine (80 mg/kg) and xylazine (20 mg/kg) and transcardially perfused with 10 mL cold aCSF followed by approximately 90–100 mL fixative containing 4% paraformaldehyde and 15% picric acid in phosphate buffer. Brains were post-fixed overnight in the same solution and sectioned at 50 μm using a Vibratome (Leica).

Sections were mounted using fluorescent mounting medium and examined using an Olympus BX51 epifluorescence microscope, Olympus VS200 slide scanner and/or Olympus FV1000 confocal microscope. Viral mCherry fluorescence within the SN was used to verify injection placement and expression.

### Reconstruction of SN viral-expression sites

For the behavioural cohort, the extent of viral expression within the SN was reconstructed across three rostrocaudal levels using the Paxinos and Franklin mouse brain atlas. Coronal sections were matched to atlas sections at approximately −2.80, −3.28 and −3.80 mm relative to bregma.

For each animal, the boundaries of mCherry-positive viral expression were manually delineated in Fiji/ImageJ and transferred to the corresponding atlas level. Individual contours were subsequently assembled and overlaid in Adobe Illustrator using 5% transparency for each mouse. Reconstructions included 12 control and 15 Cre-injected animals.

### Behavioural apparatus

Operant testing was conducted in eight Med Associates conditioning chambers (15.9 cm W × 14.0 cm H × 12.7 cm L). A food magazine equipped with infrared detectors was positioned centrally on one wall and flanked by two retractable response levers. A house light was positioned on the opposite wall, and chambers were enclosed within sound-attenuating boxes. Experimental contingencies and data acquisition were controlled using Med-PC V software.

### Food restriction, habituation and instrumental training

Food restriction commenced approximately one week before operant training and was adjusted to maintain mice at approximately 85–90% of their free-feeding body weight. Water remained available ad libitum.

Mice underwent two days of operant-chamber habituation. The first exposure lasted 15 min and the second 20 min. During habituation, food pellets were dispensed according to a random-time 60-s schedule while both levers remained retracted.

Mice were subsequently trained to lever press for 20-mg grain-based chocolate-flavoured pellets (Bio-Serv). During initial training, only one lever was available, and every lever press resulted in delivery of one pellet. Sessions terminated after 20 rewards or 20 min, whichever occurred first. Animals received two training sessions per day, one session with the left lever and one session with the right lever available.

### Fixed- and variable-ratio training

Following acquisition of lever pressing, one lever was permanently assigned to a fixed-ratio (FR) schedule and the opposite lever to a variable-ratio (VR) schedule. Under an FR5 schedule, exactly five lever presses were required to earn one reward. Under variable-ratio schedules, the response requirement varied unpredictably between rewards while averaging the stated ratio. The ratio requirement for the FR lever was maintained at 5 responses to earn an outcome throughout the experiment. The VR ratio changed depending on the experimental manipulation. VR2, VR5 and VR10 therefore required, on average, 2, 5 and 10 responses per completed reward, respectively. Lever assignments were counterbalanced across animals and remained unchanged throughout subsequent testing.

### Effort-based decision-making task

To establish a baseline of responding on each lever, mice received seven consecutive sessions of FR5-versus-VR5 training to establish stable instrumental responding and familiarity with both contingencies. Following baseline training, mice were tested in an effort-based decision-making task in which they chose between the constant FR5 option and a VR option whose effort requirement and reward magnitude were systematically manipulated. In the initial phase, both lever outcomes consisted of one food pellet. During the first three test sessions, mice chose between the FR5 lever and a low effort (VR2) option on the opposite lever. In subsequent test sessions, mice chose between the FR5 lever and an equal effort (VR5) option, or a high effort (VR10) option. Each choice condition was tested for three consecutive sessions. Following completion of the equal-reward phase, mice entered a reward–effort conflict phase. The FR5 option continued to deliver one pellet, whereas the VR option delivered three pellets. During the first three test sessions, mice chose between the FR5 and a low effort (VR2) option delivering 3 pellets. In the subsequent three test sessions, mice chose between the FR5 and a high effort (VR10) delivering 3 pellets. Sessions were capped at 40 rewards with no time limit on session duration.

### Quantification of lever preference and adaptation

The primary measure of effort allocation was the percentage of total lever presses directed to the VR option:

VR preference (%) = [VR presses / (VR presses + FR presses)] × 100.

VR preference was calculated separately for each animal and session. To quantify within-animal adaptation across repeated exposure to each contingency, an adaptation score was calculated as:

ΔVR preference = VR preference(S3) − VR preference(S1).

Positive values indicate an increase in VR responding across repeated sessions, whereas negative values indicate progressive reduction in VR preference.

Total lever presses were additionally quantified as a measure of overall instrumental output. Furthermore, initial VR preference during the first session was compared across choice conditions to determine whether subsequent group differences could be explained by pre-existing differences in lever preference.

### Trial-by-trial transition analysis

To examine the sequential organisation of choices, rewarded choice sequences were reconstructed from the raw Med Associates event arrays. Lever-identity arrays and reward-delivery arrays were aligned, and only events associated with completed reward delivery were retained. Each rewarded event was classified as resulting from either the FR or VR option.

Sequential pairs of rewarded choices were then classified as VR→VR, VR→FR, FR→VR or FR→FR transitions.

For each mouse and session, the probability of switching from VR to FR was calculated as: P(VR→FR) = N(VR→FR) / [N(VR→VR) + N(VR→FR)].

Similarly:

P(FR→VR) = N(FR→VR) / [N(FR→FR) + N(FR→VR)].

Thus, each probability was normalised to the number of eligible transition opportunities following the relevant preceding choice.

### Progressive-ratio task

Motivational persistence under progressively increasing response requirements was assessed using a progressive-ratio schedule. Animals responded on a single reinforced lever for one food pellet. For half the animals in each treatment group, the lever used for the progressive ratio test was the previously trained FR lever, and for the remaining animals it was the VR lever.

Successive response requirements followed the exponential progression:

*requirement(n) = round[5 × exp(0.2n) − 5]*

Sessions terminated after three min of inactivity or two h had elapsed. Breakpoint was defined as the highest ratio requirement successfully completed. Total lever presses and final response requirement were also quantified. For the persistence-curve representation, each animal was classified at each progressive-ratio stage according to whether it successfully reached/completed that stage.

We additionally fitted an exponential decay function to the mean response rates: *y=a*exp(x*-b)* with *y* being the response rate and *x* being the trial number. The peak response rate (*a*) and decay rate (*b*) were estimated using the non-linear least squares (nls) function in R for each animal independently and group level statistics performed on these values.

### Extinction task

Extinction of previously reinforced instrumental responding was assessed during a 30-min session in which one lever was available; however, lever pressing no longer resulted in food delivery.

Lever presses were recorded in 10-s intervals and subsequently aggregated into 5-min bins for the primary analysis. A higher-resolution analysis using 2-min bins was performed as a robustness analysis.

To quantify within-session suppression, presses emitted during the first 10 min and final 10 min of extinction were summed for each mouse. Percentage suppression was calculated as: Suppression of responding (%) = [(presses0–10 min − presses20–30 min) / presses0–10 min] × 100.

We additionally fitted an exponential decay function to the mean response rates: *y=a*exp(x*-b)* with *y* being the response rate and *x* being the trial number. The peak response rate (*a*) and decay rate (*b*) were estimated using the non-linear least squares (nls) function in R for each animal independently and group level statistics performed on these values.

Total lever presses across the complete 30-min extinction session were additionally quantified to distinguish altered extinction dynamics from differences in overall instrumental output.

### Quantification and statistical analysis

Statistical analyses were performed using GraphPad Prism 11, MATLAB R2026a, and R, and custom analysis scripts where indicated. Electrophysiological analyses additionally used AxoGraph and/or Signal. Unless otherwise indicated, data are presented as mean ± SEM, individual points represent individual recordings or animals as specified in the figure legends, all statistical tests were two-tailed and significance was set at P < 0.05.

## Supplementary Information

**Extended data Fig. 1.**
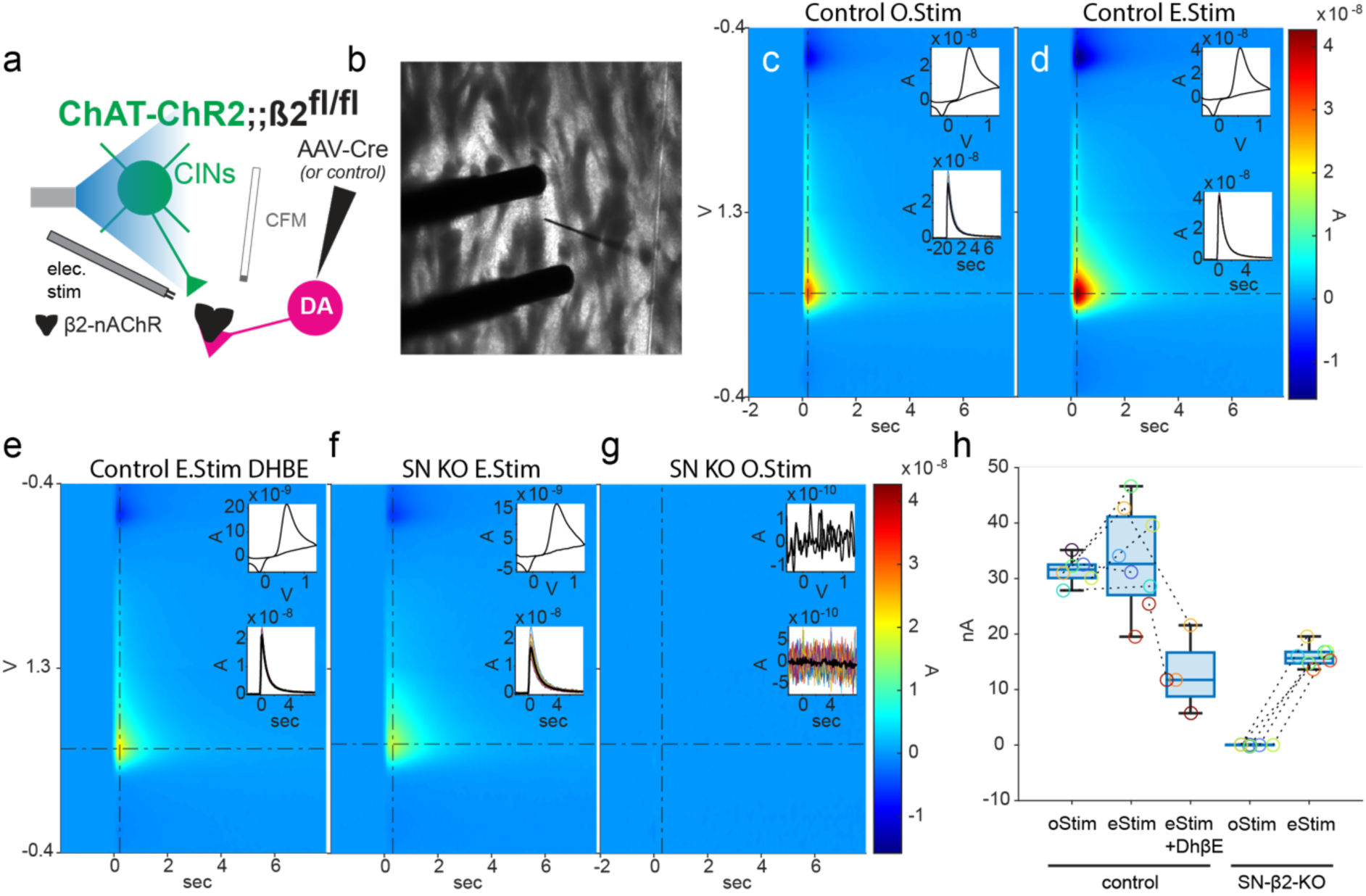
Conditional β2-nAChR deletion selectively abolishes cholinergic, but not β2-independent electrically evoked dopamine release. (a) Experimental configuration for FSCV recordings in acute striatal slices from ChAT-ChR2;Chrnb2fl/fl mice following SN injection of AAV-hSyn-Cre-mCherry or control AAV. Dopamine release was evoked by optogenetic activation of CINs or local electrical stimulation. (b) Representative DIC image showing the carbon-fibre microelectrode and bipolar stimulating electrode in dorsal striatum. (c-g) Representative FSCV pseudocolour plots with corresponding cyclic voltammograms and dopamine transients under the indicated conditions: control optical stimulation (c), control electrical stimulation (d), control electrical stimulation in DHβE (e), SN-β2 KO electrical stimulation (f) and SN-β2 KO optical stimulation (g). (h) Quantification of peak evoked dopamine concentration under the indicated stimulation conditions. Box plots show the median, interquartile range and minimum-to-maximum values; points represent individual measurements, with points of the same colour obtained from the same mouse. Dashed lines connect paired recordings obtained from the same slice.

**Extended Data Fig. 2.**
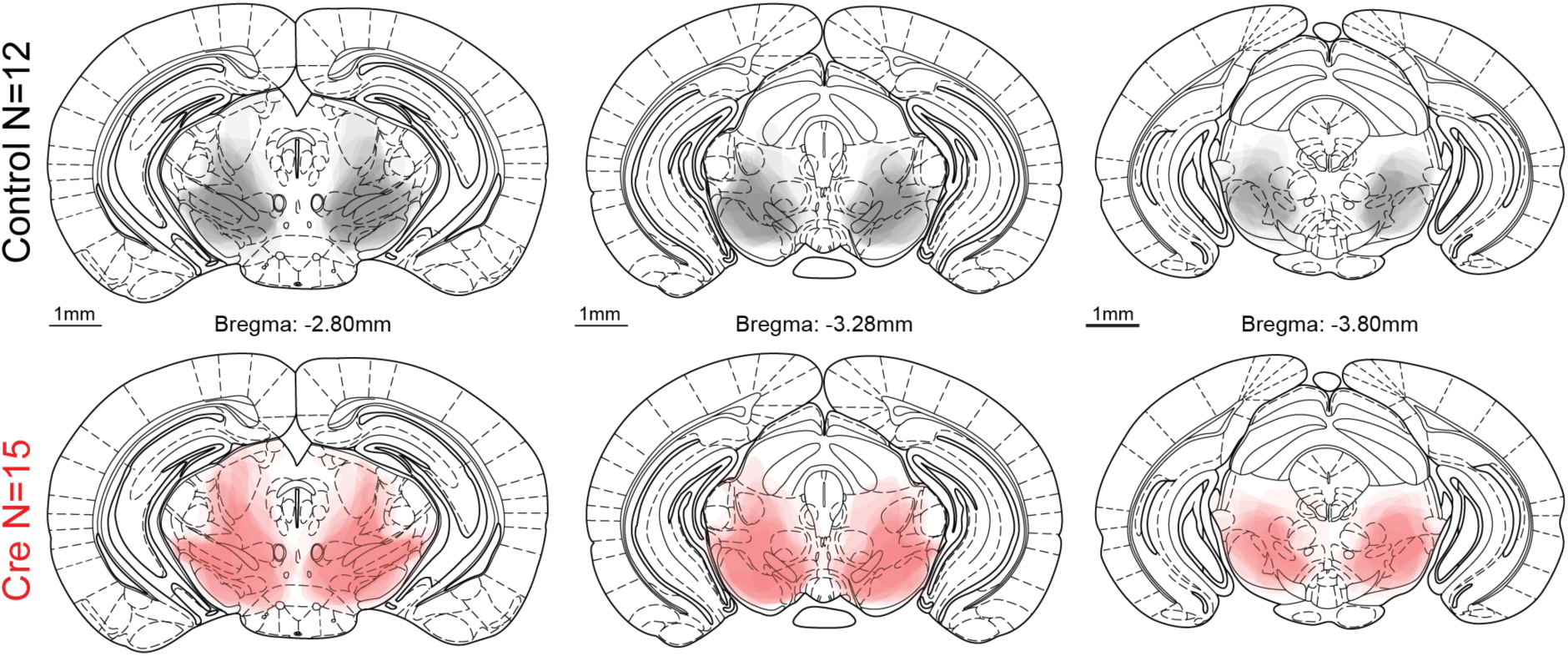
Atlas-based reconstruction of viral injection sites in the behavioural cohort. Injection contours from individual control mice (n = 12; black) and SN β2 knockout mice (n = 15; red) were overlaid at three representative rostrocaudal levels of the substantia nigra (Bregma −2.80, −3.28 and −3.80 mm) using matching coronal sections to the Paxinos and Franklin mouse brain atlas.

**Extended Data Fig. 3.**
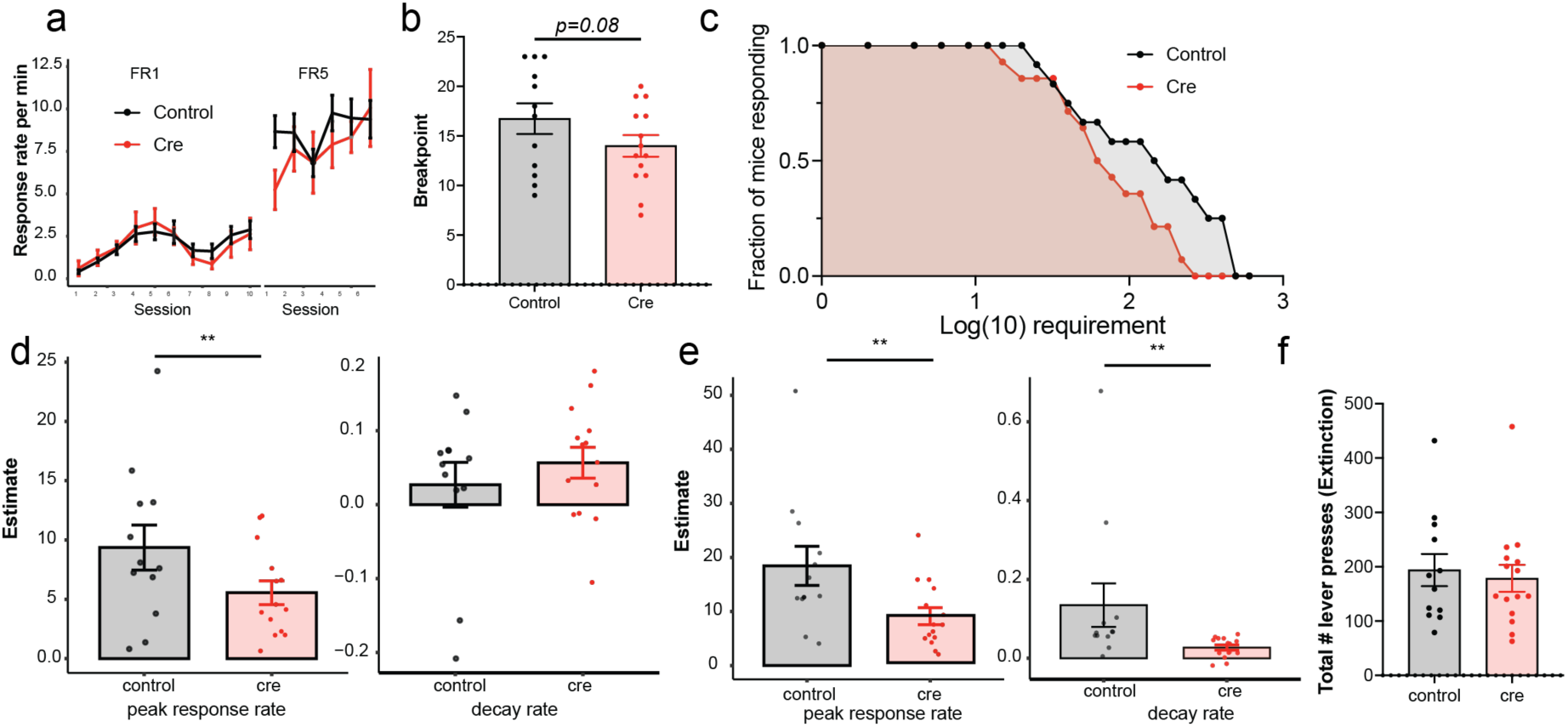
Additional analyses of progressive-ratio and extinction behaviour. (a) Acquisition of the instrumental lever-press response before progressive-ratio testing in control and SN-β2 knockout mice. (b) Breakpoint reached during the progressive-ratio session. (c) Fraction of mice remaining engaged as the response requirement increased during the progressive-ratio schedule. (d) Estimates of the peak response rate and decay rate from the progressive ratio task (e) Estimates of the peak response rate and decay rate from the extinction task (f) Total number of lever presses emitted during the 30-min extinction session. Points represent individual animals; bars show mean ± s.e.m. Statistical comparisons were performed using two-tailed unpaired tests.

**Extended Data Fig. 4.**
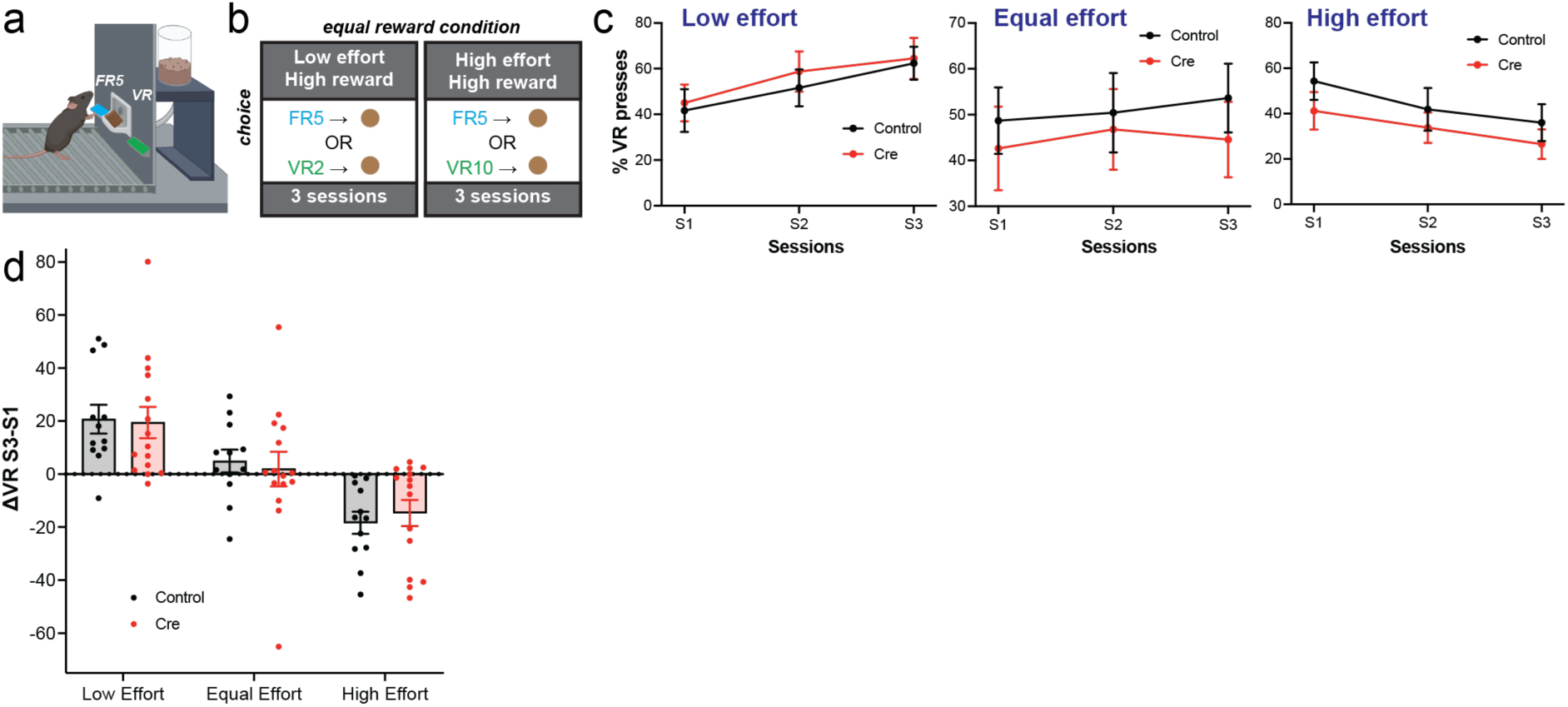
Effort-dependent choice under equal-reward conditions. (a) Schematic of the operant choice task. Mice chose between a fixed-ratio (FR5) and variable-ratio (VR) lever, with both options delivering the same reward. (b) Task contingencies. In the low-effort condition, mice chose between FR5 and VR2; in the equal-effort condition mice chose between FR5 and VR5; and in the high-effort condition, mice chose between FR5 and VR10. Reward magnitude was held constant across options. (c) Percentage of responses allocated to the VR option across three sessions under low-, equal- and high-effort conditions in control and SN-β2 knockout mice. (d) Change in VR choice from session 1 to session 3 (ΔVR S3–S1) across the three effort conditions. Points represent individual animals; bars show mean ± s.e.m.

**Extended Data Fig. 5.**
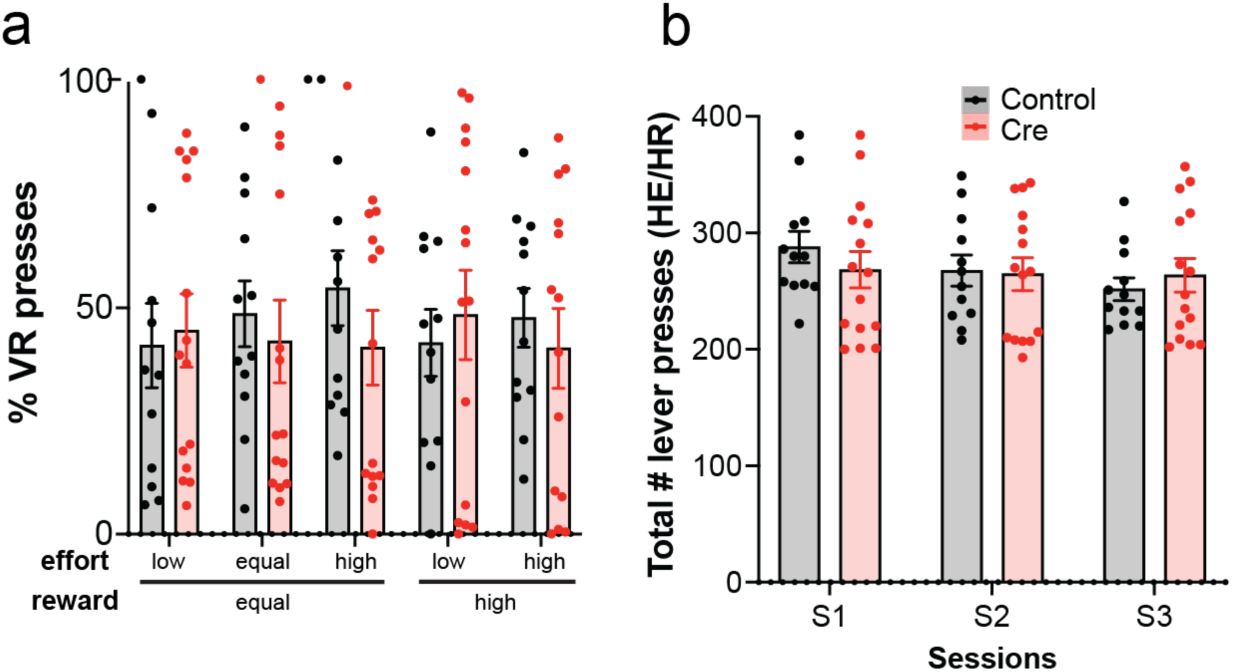
Initial choice allocation and overall responding in the effort–reward task. (a) Total number of lever presses across the three high-effort/high-reward (HE/HR) sessions in control and SN-β2 knockout mice. (b) Percentage of responses allocated to the variable-ratio (VR) option during session 1 across the different effort–reward conditions. Points represent individual animals; bars show mean ± s.e.m.

**Extended Data Fig. 6.**
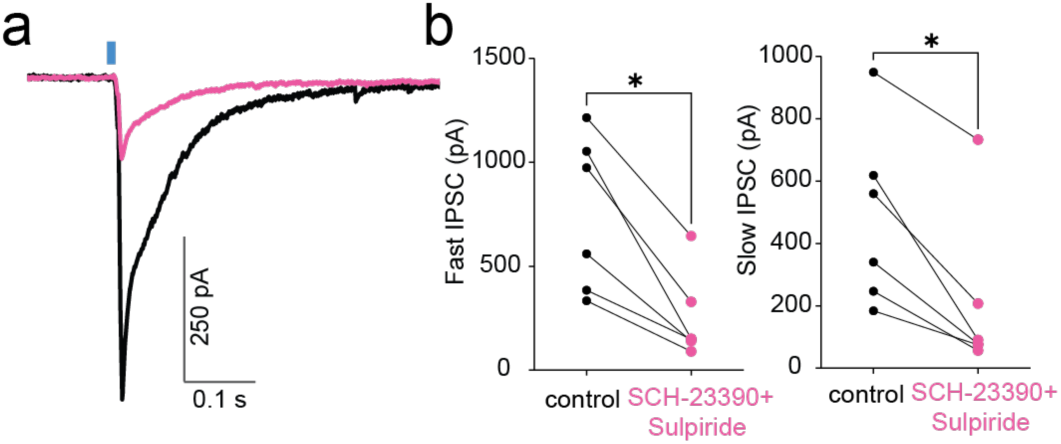
Dopamine receptor blockade reduces CIN-mediated disynaptic inhibition of SPNs. (a) Representative CIN-evoked IPSCs recorded from SPNs before and after bath application of the D1- and D2-family dopamine receptor antagonists SCH-23390 (10 μM) and sulpiride (5 μM). (b) Peak amplitude of the fast IPSC component before and after dopamine receptor blockade. (c) Peak amplitude of the slow IPSC component before and after dopamine receptor blockade. Connected points represent paired recordings from the same cell. P < 0.05, two-tailed Wilcoxon matched-pairs signed-rank test.

## References

1 . Threlfell, S., Lalic, T., Platt, N.J., Jennings, K.A., Deisseroth, K. & Cragg, S.J. Striatal dopamine release is triggered by synchronized activity in cholinergic interneurons. Neuron 75, 58–64 (2012).

2 . Cachope, R., Mateo, Y., Mathur, B.N., Irving, J., Wang, H.L., Morales, M., Lovinger, D.M. & Cheer, J.F. Selective activation of cholinergic interneurons enhances accumbal phasic dopamine release: setting the tone for reward processing. Cell Rep 2, 33–41 (2012).

3 . Kramer, P.F., Brill-Weil, S.G., Cummins, A.C., Zhang, R., Camacho-Hernandez, G.A., Newman, A.H., Eldridge, M.A.G., Averbeck, B.B. & Khaliq, Z.M. Synaptic-like axo-axonal transmission from striatal cholinergic interneurons onto dopaminergic fibers. Neuron 110, 2949–2960 e2944 (2022).

4 . Krok, A.C., Maltese, M., Mistry, P., Miao, X., Li, Y. & Tritsch, N.X. Intrinsic dopamine and acetylcholine dynamics in the striatum of mice. Nature 621, 543–549 (2023).

5 . Chantranupong, L., Beron, C.C., Zimmer, J.A., Wen, M.J., Wang, W. & Sabatini, B.L. Dopamine and glutamate regulate striatal acetylcholine in decision-making. Nature 621, 577–585 (2023).

6 . Touponse, G.C., Pomrenze, M.B., Yassine, T., Denomme, N., Wang, M., Mehta, V., Zhang, Z., Malenka, R.C. & Eshel, N. Cholinergic modulation of dopamine release drives effortful behaviour. Nature 651, 1020–1029 (2026).

7 . Salamone, J.D., Correa, M., Farrar, A. & Mingote, S.M. Effort-related functions of nucleus accumbens dopamine and associated forebrain circuits. Psychopharmacology (Berl) 191, 461–482 (2007).

8 . Niv, Y., Daw, N.D., Joel, D. & Dayan, P. Tonic dopamine: opportunity costs and the control of response vigor. Psychopharmacology (Berl) 191, 507–520 (2007).

9 . Salamone, J.D. & Correa, M. The mysterious motivational functions of mesolimbic dopamine. Neuron 76, 470–485 (2012).

10 . Salamone, J.D., Correa, M., Yohn, S., Lopez Cruz, L., San Miguel, N. & Alatorre, L. The pharmacology of effort-related choice behavior: Dopamine, depression, and individual differences. Behav Processes 127, 3–17 (2016).

11 . Salamone, J.D. & Correa, M. The Neurobiology of Activational Aspects of Motivation: Exertion of Effort, Effort-Based Decision Making, and the Role of Dopamine. Annu Rev Psychol 75, 1–32 (2024).

12 . Berke, J.D. What does dopamine mean? Nat Neurosci 21, 787–793 (2018).

13 . Hamid, A.A., Pettibone, J.R., Mabrouk, O.S., Hetrick, V.L., Schmidt, R., Vander Weele, C.M., Kennedy, R.T., Aragona, B.J. & Berke, J.D. Mesolimbic dopamine signals the value of work. Nat Neurosci 19, 117–126 (2016).

14 . Mohebi, A., Pettibone, J.R., Hamid, A.A., Wong, J.T., Vinson, L.T., Patriarchi, T., Tian, L., Kennedy, R.T. & Berke, J.D. Dissociable dopamine dynamics for learning and motivation. Nature 570, 65–70 (2019).

15 . Zhou, F.M., Liang, Y. & Dani, J.A. Endogenous nicotinic cholinergic activity regulates dopamine release in the striatum. Nat Neurosci 4, 1224–1229 (2001).

16 . Rice, M.E. & Cragg, S.J. Nicotine amplifies reward-related dopamine signals in striatum. Nat Neurosci 7, 583–584 (2004).

17 . Zhang, H. & Sulzer, D. Frequency-dependent modulation of dopamine release by nicotine. Nat Neurosci 7, 581–582 (2004).

18 . Exley, R. & Cragg, S.J. Presynaptic nicotinic receptors: a dynamic and diverse cholinergic filter of striatal dopamine neurotransmission. Br J Pharmacol 153 **Suppl 1**, S283–297 (2008).

19 . Liu, C., Cai, X., Ritzau-Jost, A., Kramer, P.F., Li, Y., Khaliq, Z.M., Hallermann, S. & Kaeser, P.S. An action potential initiation mechanism in distal axons for the control of dopamine release. Science 375, 1378–1385 (2022).

20 . Zhang, Y.F., Luan, P., Qiao, Q., He, Y., Zatka-Haas, P., Zhang, G., Lin, M.Z., Lak, A., Jing, M., Mann, E.O. & Cragg, S.J. An axonal brake on striatal dopamine output by cholinergic interneurons. Nat Neurosci 28, 783–794 (2025).

21 . Aosaki, T., Tsubokawa, H., Ishida, A., Watanabe, K., Graybiel, A.M. & Kimura, M. Responses of tonically active neurons in the primate’s striatum undergo systematic changes during behavioral sensorimotor conditioning. J Neurosci 14, 3969–3984 (1994).

22 . Morris, G., Arkadir, D., Nevet, A., Vaadia, E. & Bergman, H. Coincident but distinct messages of midbrain dopamine and striatal tonically active neurons. Neuron 43, 133– 143 (2004).

23 . Apicella, P., Legallet, E. & Trouche, E. Responses of tonically discharging neurons in the monkey striatum to primary rewards delivered during different behavioral states. Exp Brain Res 116, 456–466 (1997).

24 . Sarpong, G.A., Pass, R., Liyanagama, K., Chu, K.Y., Kurima, K., Akamine, Y., Chouinard, J.A., Looger, L.L. & Wickens, J.R. Spatially heterogeneous acetylcholine dynamics in the striatum promote behavioral flexibility. Nat Commun 16, 10877 (2025).

25 . Bradfield, L.A., Bertran-Gonzalez, J., Chieng, B. & Balleine, B.W. The thalamostriatal pathway and cholinergic control of goal-directed action: interlacing new with existing learning in the striatum. Neuron 79, 153–166 (2013).

26 . Aoki, S., Liu, A.W., Zucca, A., Zucca, S. & Wickens, J.R. Role of Striatal Cholinergic Interneurons in Set-Shifting in the Rat. J Neurosci 35, 9424–9431 (2015).

27 . Alatriste-Leon, H., Verma-Rodriguez, A.K., Ramirez-Jarquin, J.O. & Tecuapetla, F. Perturbations in the Activity of Cholinergic Interneurons in the Dorsomedial Striatum Impairs the Update of Actions to an Instrumental Contingency Change. Neuroscience 439, 287–300 (2020).

28 . Costa, K.M., Zhang, Z., Deutsch, D., Zhuo, Y., Li, G., Li, Y. & Schoenbaum, G. Dopamine and acetylcholine correlations in the nucleus accumbens depend on behavioral task states. Curr Biol 35, 1400–1407 e1403 (2025).

29 . Jang, H.J., McMahon Ward, R., Golden, C.E.M. & Constantinople, C.M. Acetylcholine demixes heterogeneous dopamine signals for learning and moving. Nat Neurosci 29, 840–850 (2026).

30 . Morgenstern, N.A., Isidro, A.F., Israely, I. & Costa, R.M. Pyramidal tract neurons drive amplification of excitatory inputs to striatum through cholinergic interneurons. Sci Adv 8, eabh4315 (2022).

31 . Matityahu, L., Hobel, Z.B., Berkowitz, N., Malgady, J.M., Gilin, N., Plotkin, J.L. & Goldberg, J.A. Synchronous activation of striatal cholinergic interneurons induces local serotonin release. Nat Commun 17 (2026).

32 . English, D.F., Ibanez-Sandoval, O., Stark, E., Tecuapetla, F., Buzsaki, G., Deisseroth, K., Tepper, J.M. & Koos, T. GABAergic circuits mediate the reinforcement-related signals of striatal cholinergic interneurons. Nat Neurosci 15, 123–130 (2012).

33 . Faust, T.W., Assous, M., Tepper, J.M. & Koos, T. Neostriatal GABAergic Interneurons Mediate Cholinergic Inhibition of Spiny Projection Neurons. J Neurosci 36, 9505–9511 (2016).

34 . Kocaturk, S., Guven, E.B., Shah, F., Tepper, J.M. & Assous, M. Cholinergic control of striatal GABAergic microcircuits. Cell Rep 41, 111531 (2022).

35. Kocaturk, S., Guven, E.B., Tepper, J.M. & Assous, M. Long-range inhibitory control of cholinergic network dynamics in the striatum. bioRxiv (2026).

36 . Witten, I.B., Lin, S.C., Brodsky, M., Prakash, R., Diester, I., Anikeeva, P., Gradinaru, V., Ramakrishnan, C. & Deisseroth, K. Cholinergic interneurons control local circuit activity and cocaine conditioning. Science 330, 1677–1681 (2010).

37 . Assous, M. Striatal cholinergic transmission. focus on nicotinic receptors’ influence in striatal circuits. Eur J Neurosci (2021).

38. Belal, M., Perez-Rosello, T., Guven, E.B., Kocaturk, S., Xie, Z., Ilijic, E., Tkatch, T., Li, J., Dauer, W., Assous, M., Tepper, J.M., Clarke, V.R.J. & Surmeier, D.J. Cholinergic interneuron control of GABAergic circuits targeting spiny projection neurons is disrupted in parkinsonian models. bioRxiv, 2026.2006.2017.732978 (2026).

39 . Nelson, A.B., Hammack, N., Yang, C.F., Shah, N.M., Seal, R.P. & Kreitzer, A.C. Striatal cholinergic interneurons Drive GABA release from dopamine terminals. Neuron 82, 63–70 (2014).

40 . El-Brolosy, M.A. & Stainier, D.Y.R. Genetic compensation: A phenomenon in search of mechanisms. PLoS Genet 13, e1006780 (2017).

41 . Wanat, M.J., Kuhnen, C.M. & Phillips, P.E. Delays conferred by escalating costs modulate dopamine release to rewards but not their predictors. J Neurosci 30, 12020– 12027 (2010).

42 . Day, J.J., Jones, J.L., Wightman, R.M. & Carelli, R.M. Phasic nucleus accumbens dopamine release encodes effort- and delay-related costs. Biol Psychiatry 68, 306– 309 (2010).

43 . Cui, G., Jun, S.B., Jin, X., Pham, M.D., Vogel, S.S., Lovinger, D.M. & Costa, R.M. Concurrent activation of striatal direct and indirect pathways during action initiation. Nature 494, 238–242 (2013).

44 . Nonomura, S., Nishizawa, K., Sakai, Y., Kawaguchi, Y., Kato, S., Uchigashima, M., Watanabe, M., Yamanaka, K., Enomoto, K., Chiken, S., Sano, H., Soma, S., Yoshida, J., Samejima, K., Ogawa, M., Kobayashi, K., Nambu, A., Isomura, Y. & Kimura, M. Monitoring and Updating of Action Selection for Goal-Directed Behavior through the Striatal Direct and Indirect Pathways. Neuron 99, 1302–1314 e1305 (2018).

45 . Peak, J., Chieng, B., Hart, G. & Balleine, B.W. Striatal direct and indirect pathway neurons differentially control the encoding and updating of goal-directed learning. Elife 9 (2020).

46. Miller-Hansen, A.J., Zhu, M., Kovaleski, R.F., Demir, B. & Lerner, T.N. Thalamus orchestrates local acetylcholine-dependent dopamine release in the learning striatum. bioRxiv (2026).

47 . Goldbach, H.C., Rimondini, R., Swanson, E.S., Shin, J.H., Authement, M.E., Anderson, L.G., Kwon, H.B., Paletzki, R., Gerfen, C.R., Amarante, L.M., Krauzlis, R.J. & Alvarez, V.A. Cholinergic-dependent dopamine signals in mouse dorsomedial striatum are regulated by frontal but not sensory cortices. Nat Commun 17 (2026).

48 . Goldberg, J.A., Ding, J.B. & Surmeier, D.J. Muscarinic modulation of striatal function and circuitry. Handb Exp Pharmacol, 223–241 (2012).

49 . Tanimura, A., Pancani, T., Lim, S.A.O., Tubert, C., Melendez, A.E., Shen, W. & Surmeier, D.J. Striatal cholinergic interneurons and Parkinson’s disease. Eur J Neurosci 47, 1148–1158 (2018).

50 . Reynolds, J.N.J., Avvisati, R., Dodson, P.D., Fisher, S.D., Oswald, M.J., Wickens, J.R. & Zhang, Y.F. Coincidence of cholinergic pauses, dopaminergic activation and depolarisation of spiny projection neurons drives synaptic plasticity in the striatum. Nat Commun 13, 1296 (2022).

